# A Lymph Node Targeted Amphiphile Vaccine Induces Potent Cellular and Humoral Immunity to SARS-CoV-2

**DOI:** 10.1101/2020.08.17.251728

**Authors:** Martin P. Steinbuck, Lochana M. Seenappa, Aniela Jakubowski, Lisa K. McNeil, Christopher M. Haqq, Peter C. DeMuth

**Affiliations:** Elicio Therapeutics, One Kendall Square, Suite 14303, Cambridge, MA 02139

## Abstract

The SARS-CoV-2 pandemic has led to public health, economic, and social consequences that mandate urgent development of effective vaccines to contain or eradicate infection. To that end, we evaluated a novel amphiphile (AMP) vaccine adjuvant, AMP-CpG, composed of diacyl lipid-modified CpG, admixed with the SARS-CoV-2 Spike-2 receptor binding domain protein as a candidate vaccine (ELI-005) in mice. AMP immunogens are efficiently delivered to lymph nodes, where innate and adaptive immune responses are generated. Compared to alum, AMP immunization induced >25-fold higher antigen-specific T cells which produced multiple Th1 cytokines and trafficked into lung parenchyma and respiratory secretions. Antibody responses favored Th1 isotypes (IgG2bc, IgG3) and potently neutralized Spike-2-ACE2 receptor binding, with titers 265-fold higher than the natural immune response from convalescent COVID-19 patients; responses were maintained despite 10-fold dose-reduction in Spike antigen. Both cellular and humoral immune responses were preserved in aged mice. These advantages merit clinical translation to SARS-CoV-2 and other protein subunit vaccines.

## Introduction

Coronavirus disease 2019 (COVID-19), caused by severe acute respiratory syndrome coronavirus 2 (SARS-CoV-2), quickly went from a public health emergency of international concern (as declared by the World Health Organization on 30 January 2020) to a pandemic (on 11 March 2020). The pandemic has resulted in worldwide social and economic consequences as well as significant health-care challenges. The severity of COVID-19 ranges from no symptoms to mild or moderate flu-like symptoms in approximately 80% of cases^1^ to serious clinical manifestations such as severe pneumonia and acute respiratory distress syndrome in up to 20% of patients^2^. Conservative estimates of 1% case fatality predict more than 40 million global deaths^3^. Unlike related respiratory betacoronaviruses, severe acute respiratory syndrome (SARS) and Middle East respiratory syndrome (MERS), the more efficient person-to-person transmission and prevalence of asymptomatic infection with SARS-CoV-2 has required public health mitigations including quarantine, contact tracing, face masks, and social distancing to reduce morbidity and mortality. Therefore, rapid development of effective vaccines and adjuvants are urgently needed.

An optimal SARS-CoV-2 vaccine should generate potent T cell immunity alongside neutralizing antibody responses. Patient recovery from SARS-CoV-2 infection without mechanical ventilation is significantly associated with elevated T cell levels^4,5,6,7^, and T cell responses without humoral responses have proven sufficient for COVID-19 resolution^8,9,10^. Conversely, death has been associated with reduced T cell numbers in COVID-19^11^, with lymphocyte subset analyses implicating deficiency in both CD3^+^CD4^+^ and CD3^+^CD8^+^ T cells^4^. Like SARS-CoV-2, lethal MERS and SARS coronavirus infections were characterized by poor or absent T cell responses^12^. However, individuals who recovered from SARS had detectable memory T cells 17 years later^13^. Further, studies have shown that SARS-CoV-2 neutralizing antibody responses rapidly decline by 3 months post onset of symptoms, highlighting the importance of T cell responses for robust and durable protection^14,15,16^. Interestingly, vaccine studies published to date have shown modest or no T cell responses to SARS-CoV-2. Optimal SARS-CoV-2 vaccines must also produce a T helper type 1 (Th1) versus type 2 (Th2) balance favoring Th1 responses. Previous SARS^17^ and MERS^18^ vaccine candidates raised preclinical safety concerns regarding exacerbating lung disease from Th2 responses or as a result of antibody-dependent enhancement (ADE) of viral entry^19^. In addition, an optimal SARS-CoV-2 vaccine must be effective across age groups, especially in the elderly. COVID-19 mortality is frequent in patients >70 years of age, coinciding with an age-related decline in immune function and comorbidities including hypertension and diabetes^20^.

In response to the urgent need for a safe and effective SARS-CoV-2 vaccine, we are evaluating a novel amphiphile (AMP) adjuvant that efficiently accumulates into lymph nodes where protective immune responses are orchestrated. Following subcutaneous injection, common immunogen components of subunit vaccines clear by either entry into the bloodstream or through lymphatic vessels; clearance route is governed by molecular weight^21^. Low molecular weight compounds (<20 kDa) can pass efficiently through the basement membrane and tight junctions of endothelial cells lining capillary vessels to enter the blood, which circulates 10-times faster than lymph flow, leading to rapid systemic distribution of these agents to immunologically irrelevant sites and to tolerizing sites such as liver^21,22^. However, larger proteins such as albumin (65 kDa) almost exclusively transit from subcutaneous tissue into lymph^22^. The AMP technology was developed to exploit this physiology using a molecular structure that conjugates immunogens (such as the vaccine adjuvant CpG DNA) to an albumin binding moiety (such as a diacyl lipid; Figure 1B), thus facilitating physical association with albumin to enable improved lymph node biodistribution (Figure 1C). The resulting conjugate is referred to as AMP-CpG. Unlike AMP-CpG, soluble CpG traffics poorly to the lymphatic system, instead entering the blood and rapidly distributing to irrelevant tissues, thus dampening the CpG adjuvant effect^23^.

**Figure 1:**
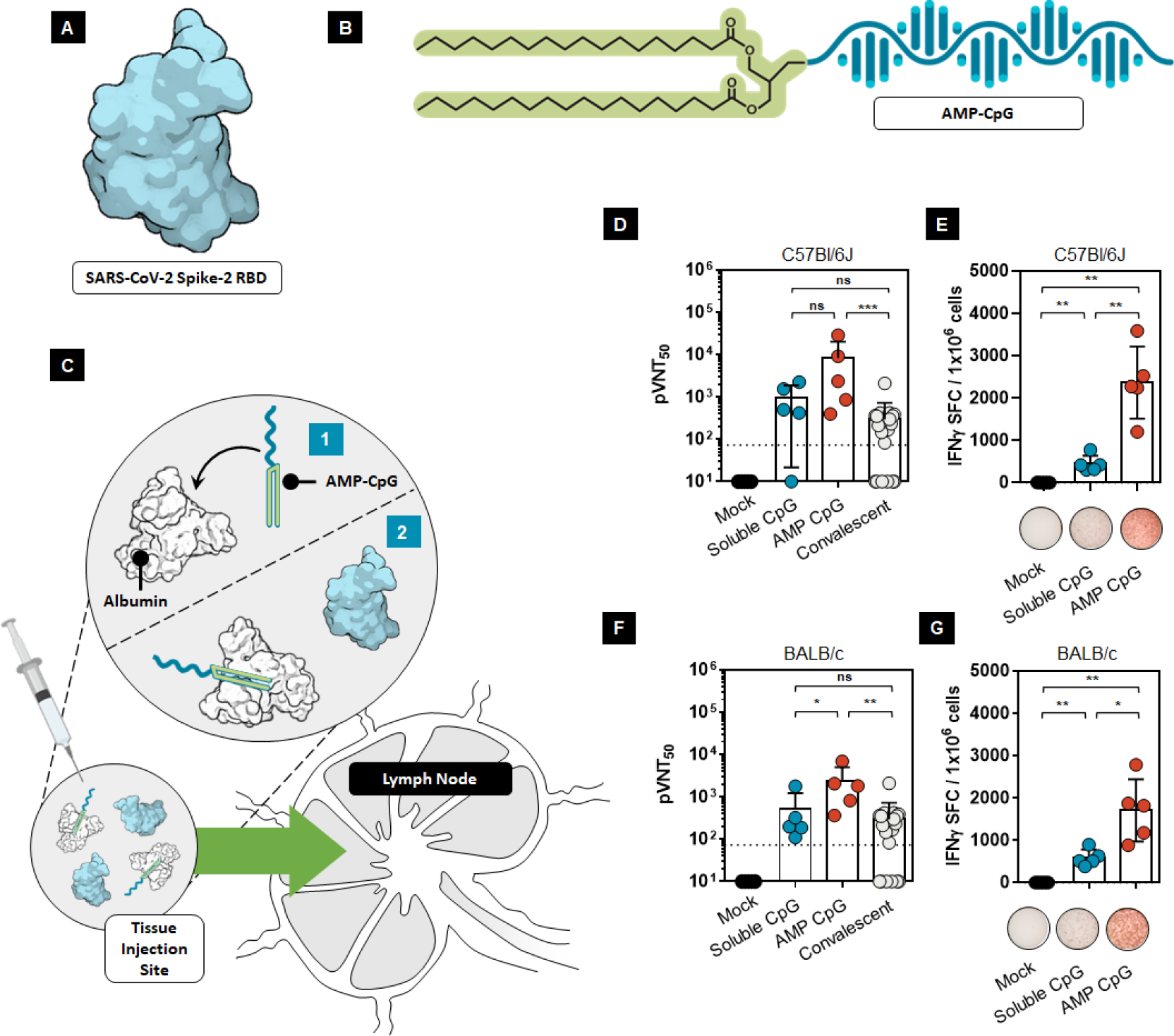
A lymph node targeted subunit vaccine elicits enhanced humoral and cellular immunity to SARS-CoV-2. Protein subunit immunization was conducted with ELI-005 consisting of **(A)** recombinant Spike-2 RBD and **(B)** AMP-CpG using a repeat dose regimen. **(C)** Mechanism of AMP-CpG targeted delivery to the lymph nodes; following subcutaneous injection, ***(1)*** AMP-CpG binds to endogenous tissue-resident albumin and ***(2)*** albumin chaperones AMP-CpG into the lymph nodes. Co-injected Spike RBD comigrates into lymph nodes. **(D-G)** C57BL/6J or BALB/C mice (n = 5 per group) were immunized on day 0, 14, 28 and 42 with 10 μg Spike RBD protein admixed with 1 nmol soluble-, or AMP-CpG, and humoral and T cell responses analyzed on day 49. Humoral and cellular immune responses were assessed in **(D-E)** C57BL/6J and **(F-G)** BALB/C mice (n = 5 per group) using **(D, F)** pseudovirus neutralization assays and **(E, G)** splenocyte IFNγ ELISpot assays, respectively. Pseudovirus neutralization activity was compared to sera/plasma from a cohort of 22 convalescent patients who had recovered from SARS-CoV-2 infection. Values depicted are mean ± standard deviation. Not detected values are shown on the baseline; * P < 0.05; ** P < 0.01; *** P < 0.001; **** P < 0.0001 by two-sided Mann-Whitney test. Pseudovirus LOD (indicated by the dotted line) was determined as mean + 90% CI calculated for mock treatment.

Previously, the diacyl lipid component of the AMP vaccine was shown to bind to albumin in subcutaneous tissue, taking advantage of albumin’s natural physiological function to transport lipids, and allowed the AMP vaccine to “hitchhike” on albumin into lymph nodes. AMP vaccines are efficiently carried into the lymph and then draining lymph nodes, where they accumulate in key antigen-presenting cells, leading to a 30- to 50-fold increase in T cell responses and antibody responses to whole recombinant protein or peptide vaccines in mice^24,25^.

The AMP technology is currently being developed for therapeutic cancer vaccines and immunotherapies that make use of the AMP strategy to enhance lymphatic delivery of antigenic peptides^26^, adjuvants, CAR-T activators for hematological and solid tumors^27^, cytokines, and immunomodulatory agents. Based on the preclinical work to support the oncology indication and the work presented in this communication, we hypothesize that AMP-CpG will combine with SARS-CoV-2 antigens to provide a safe and effective SARS-CoV-2 vaccine.

In this communication, we describe the immunogenicity of AMP-CpG paired with SARS-CoV-2 Spike-2 receptor binding domain protein (hereafter referred to as Spike RBD), which is of sufficient size (approximately 34 kDa) to predominantly traffic to lymph nodes after subcutaneous injection^21^. The coronavirus Spike protein is the largest surface virion protein and the target of neutralizing antibodies which block the interaction between SARS-CoV-2 and its angiotensin converting enzyme 2 (ACE2) human receptor^28^. Several groups have shown that Spike-based antigens serve as targets for neutralizing humoral responses to SARS^17^, MERS^18^, and SARS-CoV-2 in mice, rats, primates, and humans^29,30,31,32^. Likewise, Spike antigens contain T cell epitopes, generating cell mediated immunity in SARS^33^, MERS^12^, and SARS-CoV-2^7,34,35^.

We evaluated T-cell immunity in splenocytes, peripheral blood, cells from perfused lung tissues, and bronchoalveolar lavage (BAL) fluid. We also measured humoral responses specific to Spike RBD through enzyme-linked immunosorbent assays (ELISA), pseudovirus and surrogate neutralization assays. Further, we compared immune responses following administration of a range of antigen concentrations, and we evaluated immune responses in aged mice. The data show that immunization against Spike RBD with AMP-CpG elicits Th-1 biased cellular (with polyfunctional T cells) and humoral responses in young and aged mice. The AMP-CpG vaccine (ELI-005) could offer an optimal vaccine for general clinical use.

## Results

### Formulation Design

For assessment of the AMP technology as a possible vaccine for SARS-CoV-2, we evaluated AMP-CpG (consisting of a diacyl lipid conjugated to CpG DNA) admixed with the Spike RBD protein (the admixture of AMP-CpG and Spike RBD is referred to as ELI-005) in a repeat dose immunization regimen (Figure 1A and 1B). Following subcutaneous injection, AMP-CpG binds to endogenous tissue-resident albumin, which then travels to lymph nodes where it effectively colocalizes with lymph node resident immune cells (Figure 1C). Initial assessments in C57BL/6J and BALB/C mice receiving immunization containing AMP-CpG produced a 8- to 28-fold higher pseudovirus neutralizing titer than natural antibody responses present in human convalescent serum (obtained from recovered COVID-19 patients; Figure 1D and 1F), indicating the potential for AMP-CpG to produce neutralizing antibody responses more potent than natural immunity. By comparison, animals immunized with a dose-matched regimen containing unmodified (soluble) CpG produced neutralizing titers comparable to those observed in human convalescing patients. The results of splenocyte ELISpot assays showed that compared with soluble CpG, mice immunized with Spike RBD protein admixed with AMP-CpG elicited approximately 4-fold greater frequencies of antigen-specific functional T cells, producing IFNγ upon stimulation with Spike-derived overlapping peptides (Figure 1E and 1G).

### Cellular Immune Response

#### Splenocytes and Peripheral Blood

To further characterize the contribution of a lymph node targeted adjuvant on Spike-RBD-specific immunity we compared AMP-CpG to dose-matched formulations of antigen admixed with either soluble CpG (representative of an adjuvant with poor lymph node uptake) or alum (as a benchmark injection site depot-forming formulation). We assessed cytokine-producing cells in splenocytes and peripheral blood from C57BL/6J mice on day 35. Mice immunized with AMP-CpG had substantially higher IFNγ spot forming cells than mice dosed with soluble CpG, alum, or mock (all admixed with Spike RBD protein; Figure 2A). Approximately 43% of CD8^+^ T cells derived from peripheral blood in AMP-CpG immunized mice were cytokine producing (IFNγ, TNFα, or double-positive T cells); in comparison, approximately 13% and <2% of CD8^+^ T cells were cytokine-producing for soluble CpG-immunized mice and alum-immunized mice, respectively (Figure 2B). A similar trend was observed for CD4^+^ T cells, though percentages were relatively smaller: approximately 1.5% of T cells in peripheral blood from AMP-CpG immunized mice were cytokine-producing compared with <1% in CpG-immunized mice and <0.5% for alum-immunized mice and mock-immunized mice (Figure 2C).

**Figure 2:**
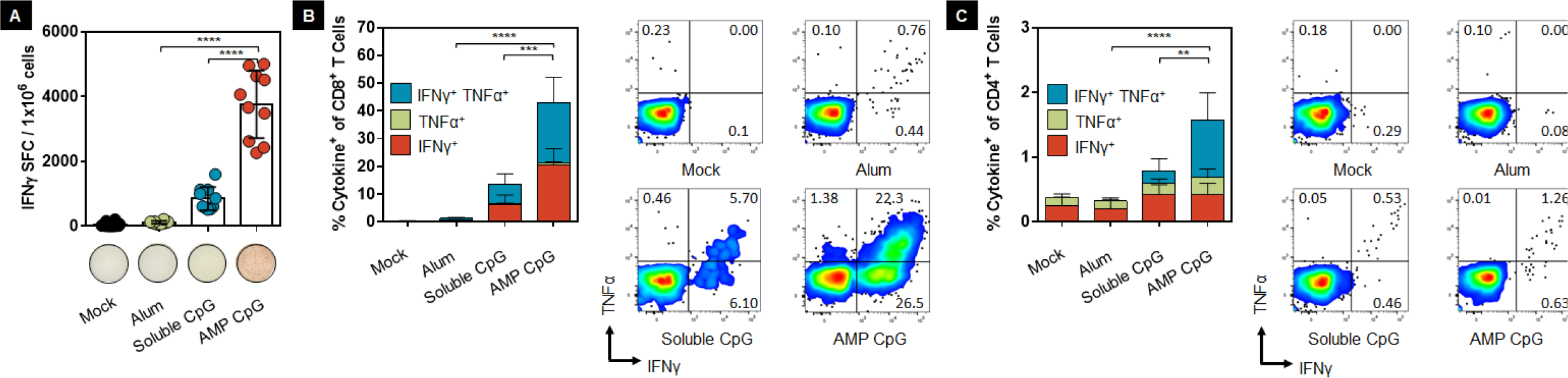
Vaccination with AMP-CpG elicits potent Spike RBD-specific CD8 and CD4 T cells in spleen and blood. C57BL/6J mice (n = 10 per group) were immunized on day 0, 14, and 28 with 10 μg Spike RBD protein admixed with 100 μg Alum, 1 nmol soluble-, or AMP-CpG, and T cell responses analyzed on day 35. **(A)** Splenocytes were stimulated with overlapping Spike peptides and assayed for IFNγ production by ELISpot assay. Shown are representative images of ELISpot and frequency of IFNγ spot forming cells (SFC) per 1×10^6^ splenocytes. **(B-C)** Peripheral blood cells were stimulated with overlapping Spike peptides and assayed by flow cytometry for intracellular cytokine production to detect antigen-specific T cell responses. Shown are frequencies of IFNγ, TNFα, and double-positive T cells among **(B)** CD8^+^ and **(C)** CD4^+^ T cells, with corresponding representative flow cytometry plots. n = 10 mice per group. Values depicted are mean ± standard deviation. * P < 0.05; ** P < 0.01; *** P < 0.001; **** P < 0.0001 by two-sided Mann-Whitney test applied to cytokine^+^ T cell frequencies.

#### Perfused Lung Tissue

To determine whether immunization could induce tissue resident T cell responses at a site of likely SARS-CoV-2 exposure, we evaluated the proportion of cytokine-producing cells from perfused lung tissue from C57BL/6J mice on day 35. As observed with responses assayed in peripheral blood, the proportion of cytokine producing cells among CD8^+^ or CD4^+^ T cells was highest in AMP-CpG-immunized mice compared with soluble CpG, alum, or mock treatment (Figure 3A and 3B). Interestingly, T cells in the lung tissue had a greater proportion of the cytokine-producing cells than observed in peripheral blood. Observations in AMP-CpG immunized mice showed that approximately 73% of CD8^+^ T cells from perfused lung tissues were cytokine-producing, with approximately 40% exhibiting polyfunctional secretion of both Th1 cytokines IFNγ and TNFα. By comparison, immunization with soluble CpG or alum induced >5-fold and >25-fold lower responses, respectively. Similar assessment of CD4^+^ T cells showed that only AMP-CpG immunized animals generated responses above background, with approximately 6% of CD4^+^ T cells producing IFNγ and/or TNFα, again exhibiting strong polyfunctional effector functionality, with the majority of these cells able to produce both IFNγ and TNFα upon antigen stimulation. These results show that the more potent lymph node action of AMP-CpG induces enhanced expansion of antigen-specific T cells with potentially beneficial tissue homing properties, establishing protective tissue resident cells at a primary site of initial viral exposure.

**Figure 3:**
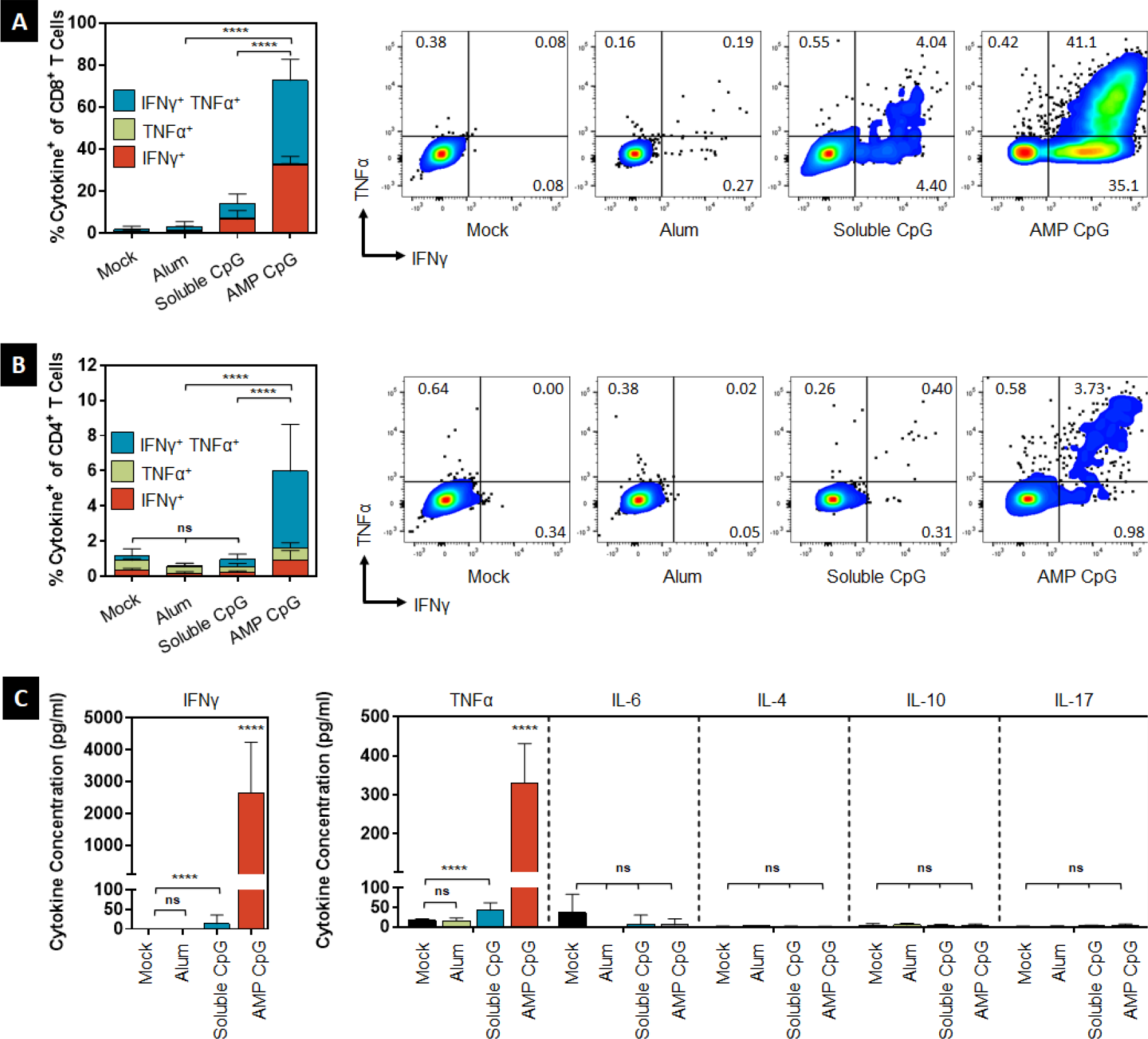
Vaccination with AMP-CpG elicits potent lung resident Spike RBD-specific CD8 and CD4 T cells. C57BL/6J mice (n = 10 per group) were immunized on day 0, 14, and 28 with 10 μg Spike-2 RBD protein admixed with 100 μg Alum, 1 nmol soluble-, or AMP-CpG, and T cell responses analyzed on day 35. **(A)** Cells collected from perfused lung tissue were stimulated with overlapping Spike peptides and assayed for cytokine production to detect antigen-specific T cell responses. Shown are frequencies of IFNγ, TNFα, and double-positive T cells among **(A)** CD8^+^ and **(B)** CD4^+^ T cells, with corresponding representative flow cytometry plots. **(C)** Cytokine concentration in supernatants from stimulated lung cells. n = 10 mice per group. Values depicted are mean ± standard deviation. * P < 0.05; ** P < 0.01; *** P < 0.001; **** P < 0.0001 by two-sided Mann-Whitney test applied to cytokine^+^ T cell frequencies or cytokine concentrations.

To more comprehensively understand the Th1/Th2/Th17 profile of the elicited T cell responses, we used a multiplexed cytokine assay to assess various cytokine concentrations from supernatants of cells collected from perfused lungs following stimulation with Spike-derived overlapping peptides. AMP-CpG immunized mice exhibited a Th1 effector profile consistent with prior assessment by flow cytometry with IFNγ and TNFα concentrations that were significantly higher than cohorts immunized with the other adjuvants (soluble CpG, alum) or mock: the IFNγ concentration was at least 200-fold higher than observed with the other adjuvants or mock, and the TNFα concentration was at least 7-fold higher than the other adjuvants or mock (Figure 3C). Concentrations of common Th2 or Th17 associated cytokines IL-4, IL-6, IL-10, and IL-17 were undetectable for all cohorts. These results further demonstrate the greatly enhanced potency and Th1-bias in T cells elicited through immunization with AMP-CpG compared with either soluble CpG or alum.

#### Bronchoalveolar (BAL) Fluid

Together with T cells in the lung parenchyma, BAL-resident T cells may be ideally positioned to rapidly respond to prevent or control nascent viral infection. To further evaluate whether lung-resident T cell responses induced by immunization could localize into lung secretions we collected BAL fluid from C57BL/6J immunized mice on day 35 and determined T cell numbers and phenotypic characteristics. We found significantly more CD8^+^ T cells in BAL fluid of AMP-CpG immunized mice than other treatment groups (Figure 4A). In addition, a significantly lower proportion of cells detected in the BAL collected from AMP-CpG immunized animals exhibited a naïve phenotype (CD44^-^, CD62L^+^; Figure 4B) with a corresponding increase in the frequency of effector memory phenotype (T_EM_; CD44^+^, CD62L^-^; Figure 4C). The CD4^+^ T cell count was enhanced relative to mock treatment and generally similar across all treatment groups (Figure 4D), but the AMP-CpG cohort showed evidence that a significantly greater proportion of the BAL-resident CD4^+^ T cells had differentiated from naïve to T_EM_ phenotype than in the other treatment groups (Figure 4F). The improved numbers and phenotype of BAL-resident T cells present in AMP-CpG immunized animals demonstrate a greater potential for early immunological detection and control at the point of viral exposure.

**Figure 4:**
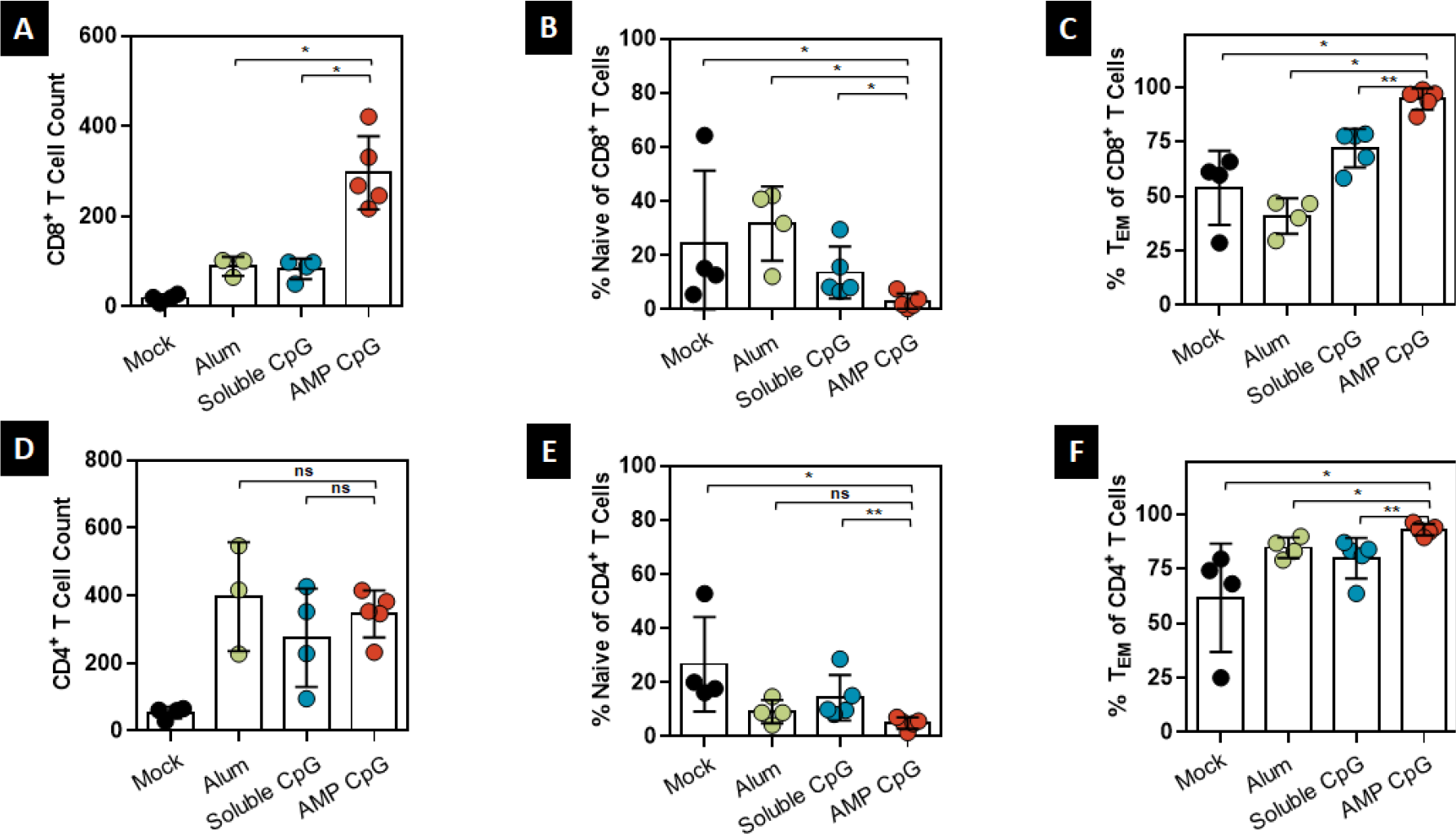
Vaccination with AMP-CpG elicits enhanced Spike RBD-specific T cell responses in BAL. C57BL/6J mice (n = 10 per group) were immunized on day 0, 14, and 28 with 10 μg Spike RBD protein admixed with 100 μg Alum, 1 nmol soluble-, or AMP-CpG, and T cell responses analyzed on day 35. Cells collected from bronchoalveolar lavage (BAL) were analyzed for T cell phenotype by flow cytometry. Shown are total count of **(A)** CD8^+^ T cells and frequencies of **(B)** naïve (CD44^-^CD62L^+^), and **(C)** effector memory (T_EM_; CD44^+^ CD62L^+^) T cells among CD8^+^ T cells. Corresponding analyses are shown for **(D)** CD4^+^ T cells, **(E)** naïve (CD44^-^CD62L^+^), and **(F)** effector memory (T_EM_; CD44^+^ CD62L^+^) T cells among CD4^+^ T cells. n = 10 mice per group. Values depicted are mean ± standard deviation. * P < 0.05; ** P < 0.01; *** P < 0.001; **** P < 0.0001 by two-sided Mann-Whitney test applied to T cell frequencies.

### Humoral Responses

#### Neutralizing Antibody

Neutralizing antibody responses to Spike RBD are generated in convalescing patients and are a primary goal for vaccines that would prevent infection or limit its severity. As in prior studies^31,36^, we assessed neutralizing antibody activity through measurement of inhibition of the Spike-RBD-ACE2 interaction in an ELISA-based surrogate assay. Results for serum collected on day 35 for cohorts of immunized C57BL/6J mice are shown in Figure 5A. Comparable levels of neutralizing activity were induced in animals immunized with AMP-CpG, soluble CpG, and alum. Comparison with samples obtained from a cohort of convalescent humans showed that the vaccine-induced responses were significantly higher than those generated through response to natural infection.

**Figure 5:**
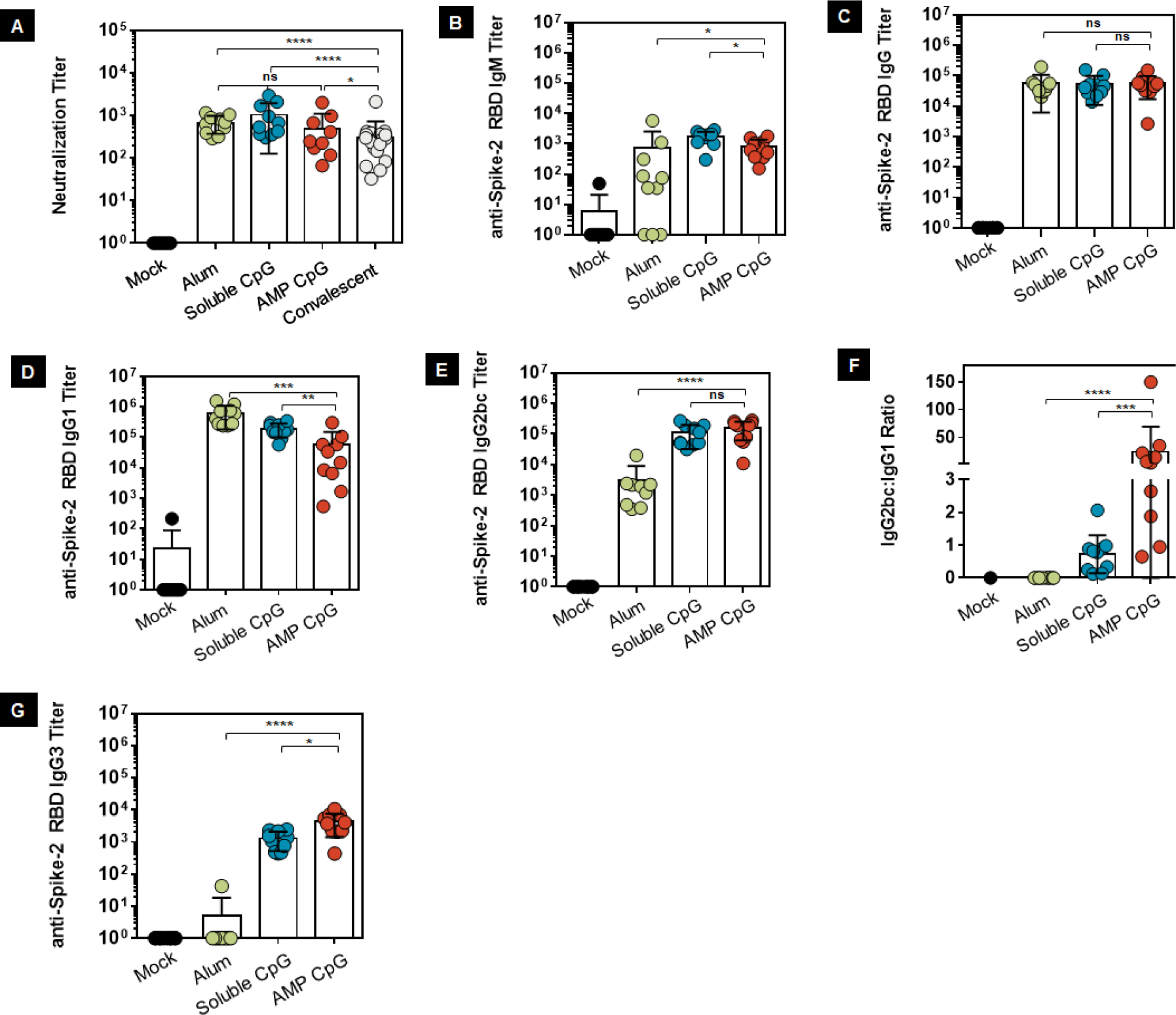
Vaccination with AMP-CpG elicits potent Th1-biased Spike RBD humoral responses. C57Bl/6 mice were immunized on days 0, 14, and 28 with 10 μg Spike RBD protein admixed with 100 μg Alum, 1 nmol soluble-, or AMP-CpG, and humoral responses analyzed on day 7 or day 35. Humoral responses specific to Spike RBD were assessed in serum from immunized animals by pseudovirus neutralization or ELISA assay. Shown are **(A)** surrogate neutralization endpoint titers on day 35, **(B-G)** endpoint titers, and endpoint titer ratios determined by ELISA for **(B)** IgM on day 7, **(C)** IgG, **(D)** IgG1, **(E)** and IgG2bc on day 35, **(F)** IgG2bc:IgG1 ratio, and **(G)** IgG3 on day 35. Pseudovirus neutralization activity was compared to sera/plasma from a cohort of 22 convalescent patients who had recovered from SARS-CoV-2 infection. n = 10 mice per group. Values depicted are mean ± standard deviation. Not detected values are shown on the baseline; * P < 0.05; ** P < 0.01; *** P < 0.001; **** P < 0.0001 by two-sided Mann-Whitney test.

#### IgM/IgG

Seven days after the initial immunization, all cohorts, except the control receiving mock immunization, showed robust Spike RBD-specific IgM responses (Figure 5B); which underwent isotype switching to produce IgG responses with similar titer following subsequent boosting immunization (Figure 5C).

#### IgG Subclasses

To assess Th1/Th2-bias in the Spike-RBD-specific IgG response elicited through immunization, we evaluated the IgG subclasses present and found that mice immunized with AMP-CpG or soluble CpG had significantly lower Th2 associated IgG1 titers (approximately 3- to 10-fold) than mice immunized with alum (Figure 5D). The reverse was true for Th1 associated IgG2bc: titers were significantly higher (approximately 50-fold) for mice immunized with AMP-CpG or soluble CpG (Figure 5E). The ratio of IgG2bc:IgG1 titer indicated a strong bias towards Th1 for AMP-CpG immunized animals, while soluble CpG and alum produced a balanced Th1/Th2 profile or Th2-dominant response, respectively (Figure 5F). Further analysis showed AMP-CpG immunized animals produced significantly higher IgG3 titers than either soluble CpG (approximately 3-fold) or alum (>800-fold) treatment groups consistent with the observed Th1-bias resulting from AMP-CpG immunization (Figure 5G).

#### Antigen Dose Sparing

Given the need for immunization against SARS-CoV-2 on a global scale, and the observed potency of AMP-CpG for induction of potent neutralizing antibodies with CD8^+^ and CD4^+^ T cells in peripheral blood, lung, and BAL, we sought to determine whether dose sparing could produce similar immune responses with reduced doses of Spike RBD antigen. Here we studied the immune response generated by repeat-dose immunization with AMP-CpG admixed with three different dose levels of Spike RBD (1, 5, and 10 μg) and compared these to responses induced by immunization with soluble CpG and alum admixed with 10 μg of antigen.

#### Cellular Immune Response

We evaluated T cell responses on day 35 in spleen, peripheral blood, and lung tissues. The results showed that the number of IFNγ-producing cells in splenocytes collected from AMP-CpG immunized C57BL/6J mice tended to increase with antigen concentration, but, even at the lowest antigen dose admixed with AMP-CpG, the number of IFNγ-producing cells was significantly higher than observed in cohorts that received the highest antigen dose (10 μg) with either soluble CpG (approximately 6-fold) or alum (>40-fold) (Figure 6A). In both peripheral blood (Figure 6B and 6C) and lung tissue (Figure 6D and 6E), the percent of CD8^+^ and CD4^+^ T cells producing cytokine was significantly higher for AMP-CpG treated mice at any concentration of antigen compared with the other adjuvants tested. Notably, no significant decrease in the frequency of cytokine-producing CD8^+^ or CD4^+^ T cells was observed in the peripheral blood of animals immunized with AMP-CpG admixed with antigen at 10, 5, or 1 µg dose levels as these were maintained at approximately 40%-50% of CD8^+^ and 2%-4% of CD4^+^ T cells. While a decreasing trend was observed in the frequency of lung-resident cytokine-producing CD8^+^ T cells in AMP-CpG immunized animals, even the 1 µg dose level produced frequencies >3-fold or >30-fold higher than animals immunized with soluble CpG or alum, respectively. This indicates a significant potential for AMP-CpG to enable at least 10-fold dose sparing of RBD protein.

**Figure 6:**
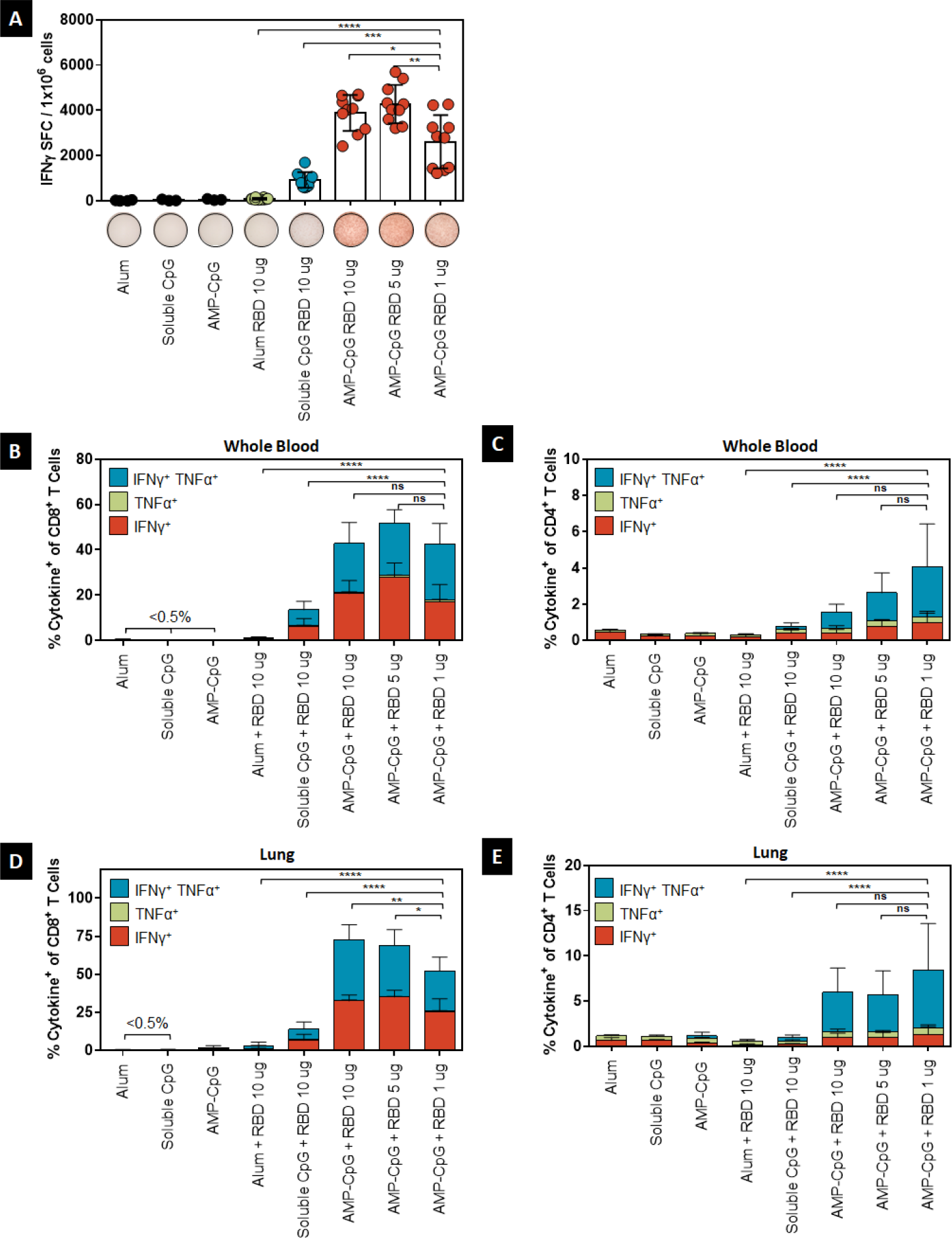
Vaccination with AMP-CpG enables dose sparing of Spike RBD to elicit cellular immunity. C57BL/6J mice were immunized on day 0, 14, and 28 with 10 μg Spike RBD protein admixed with 100 μg Alum or 1 nmol soluble-CpG. Comparator animals were dosed with 1, 5, or 10 μg Spike RBD admixed with 1 nmol AMP-CpG. Control animals were dosed with adjuvant alone. T cell responses were analyzed on day 35. **(A)** Splenocytes were stimulated with overlapping Spike peptides and assayed for IFNγ production by ELISpot assay. Shown are representative images of ELISpot and frequency of IFNγ spot forming cells (SFC) per 1×10^6^ splenocytes. Cells collected from **(B-C)** peripheral blood and **(D-E)** perfused lung tissue were stimulated with overlapping Spike peptides and assayed for intracellular cytokine production to detect antigen-specific T cell responses. Shown are frequencies of IFNγ, TNFα, and double-positive T cells among **(B)** peripheral blood CD8^+^ and **(C)** CD4^+^ T cells, and **(D)** lung CD8^+^ and **(E)** CD4^+^ T cells. n = 10 mice per group. Values depicted are mean ± standard deviation. * P < 0.05; ** P < 0.01; *** P < 0.001; **** P < 0.0001 by two-sided Mann-Whitney test applied to cytokine^+^ T cell frequencies.

#### Humoral Immune Response

On day 35 following repeat dose immunization, we assessed the induction of Spike RBD-specific antibody responses among the AMP-CpG immunized animals at each specified Spike RBD dose level for comparison to responses generated by immunization with either soluble CpG or alum at the 10 μg dose. Neutralizing activity was assessed through measurement of pseudovirus neutralization titers at the half-maximal inhibitory dilution (pVNT_50_). Similar levels of pseudovirus neutralization titers were observed for all treatment groups, at levels that were 265, 184, or 94-fold greater than those observed in convalescent human samples, for AMP-CpG, soluble CpG, and alum immunized mice, respectively (Figure 7A). Notably, these levels were maintained in animals immunized with AMP-CpG at lower Spike RBD dose levels with mean pVNT_50_ at least 115-fold greater than those measured in recovering COVID-19 patients (Figure 7B).

**Figure 7:**
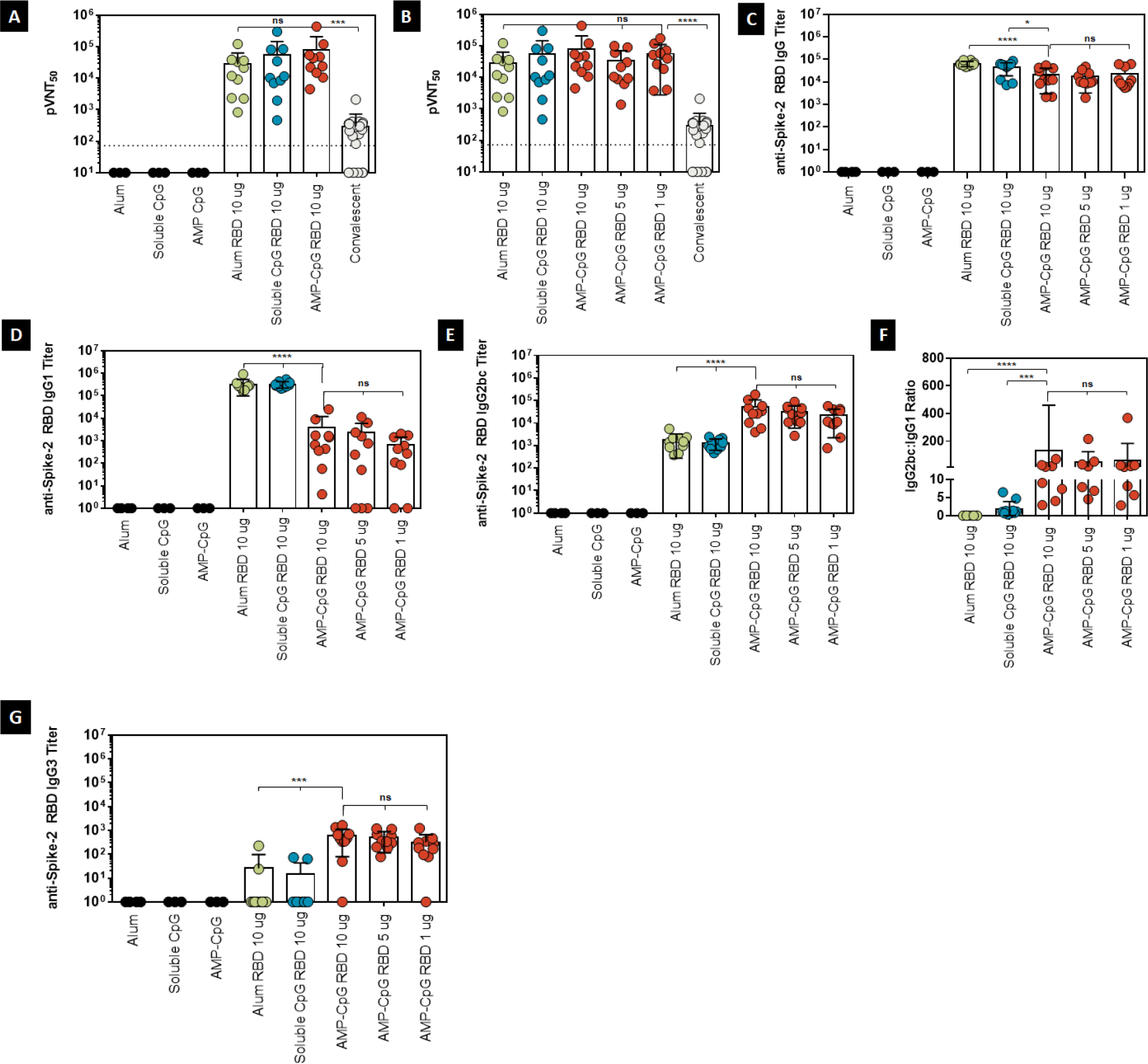
Vaccination with AMP-CpG enables dose sparing of Spike RBD to elicit humoral immunity. C57BL/6J mice were immunized on day 0, 14, and 28 with 10 μg Spike RBD protein admixed with 100 μg Alum or 1 nmol soluble-CpG. Comparator animals were dosed with 1, 5, or 10 μg Spike RBD admixed with 1 nmol AMP-CpG. Control animals were dosed with adjuvant alone. Humoral responses were analyzed on day 35. Humoral responses specific to Spike RBD were assessed in serum from immunized animals by ELISA or pseudovirus neutralization assay. Shown are **(A, B)** pseudovirus neutralization ID_50_ (pVNT_50_) on day 35, **(C-G)** endpoint titers, and endpoint titer ratios determined for **(C)** IgG, **(D)** IgG1, **(E)** IgG2bc, **(F)** IgG2bc:IgG1 ratio, and **(G)** IgG3 on day 35. Pseudovirus neutralization activity was compared to a cohort of 22 convalescent patients who had recovered from SARS-CoV-2 infection. n = 10 mice per group. Values depicted are mean ± standard deviation. Not detected values are shown on the baseline; * P < 0.05; ** P < 0.01; *** P < 0.001; **** P < 0.0001 by two-sided Mann-Whitney test. Pseudovirus LOD (indicated by the dotted line) was determined as mean + 90% CI calculated for mock treatment.

Total IgG titers were similar among the groups administered AMP-CpG, and these were reduced approximately 2-fold in comparison to titers measured among groups dosed with either soluble CpG or alum adjuvanted vaccines (Figure 7C). Isotype analysis demonstrated similar trends to those initially observed in comparison at the 10 μg dose level (Figure 5C-5G). Alum and soluble CpG immunization produced significantly higher Th2-associated IgG1 titers (approximately 100-fold) compared to all Spike RBD dose levels admixed with AMP-CpG (Figure 7D). Th1-associated IgG2bc levels were elevated approximately 20-fold in all AMP-CpG immunized animals compared with soluble CpG and alum immunized groups, with no significant difference observed with reduced Spike RBD dose level. These trends were further evident in the comparison of IgG2bc:IgG1 titer ratio (Figure 7F), where AMP-CpG containing regimens induced highly Th1-dominant isotype profile (IgG2bc:IgG1 >40), compared with more balanced and Th2-skewed responses in soluble CpG (IgG2bc:IgG1 approximately 2) and alum (IgG2bc:IgG1 <1) vaccinated animals respectively. Finally, only animals immunized with AMP-CpG showed evidence of Spike RBD-specific IgG3 titers, with comparable levels detected among all Spike RBD dose levels (approximately 500-fold over background). Further analysis showed AMP-CpG immunized animals produced significantly higher IgG3 titers than either soluble CpG (approximately 40-fold) or alum (>20-fold) treatment groups consistent with the observed Th1-bias resulting from AMP-CpG immunization (Figure 7G). Together these data demonstrate the potential for AMP-CpG to enable at least 10-fold dose sparing of Spike RBD antigen for induction of neutralizing, high titer, and optimal Th1 profile antibody responses against Spike RBD. While soluble CpG and alum induced marginally higher total IgG responses, these did not result in significant differences in neutralizing activity compared to AMP-CpG immunization. Alum and, to a lesser degree, soluble CpG responses were dominated by the Th2-associated IgG1 isotype raising the potential for a risk of toxicity in human translation based on prior outcomes in SARS and MERS vaccine development.

### Immune Response in Aged Mice

Protective immunization in at-risk populations is of utmost importance. Specifically, COVID-19 has demonstrated a higher incidence of severe disease and mortality among elderly human patients coincident with a general decline in immune function associated with aging^37,38,39^. To assess the potential of AMP-CpG to elicit effective immunity in the setting of deficient baseline immune function, we immunized aged mice to compare immune responses generated from vaccines containing AMP-CpG, soluble CpG, and alum. We further investigated the effect of Spike RBD protein dose in this model to determine the potential for AMP-CpG to enable antigen dose sparing similar to that observed in studies of young healthy mice.

#### Cellular Immune Response

We evaluated T cell responses in aged mice after repeat dose immunization with comparator vaccines on days 21 and 35. Assessment on day 21 of cytokine-producing CD8^+^ T cells in peripheral blood following Spike-derived overlapping peptide stimulation showed that AMP-CpG induced potent responses (approximately 15% of CD8^+^ T cells), greatly outperforming soluble CpG (approximately 2.5% of CD8^+^ T cells), and alum (<0.5% of CD8^+^ T cells) (Figure 8A). Although these responses were reduced approximately 2-fold compared to those observed in young healthy mice, they nonetheless exceeded those generated by the soluble CpG and alum comparators by 6- and 30-fold, respectively. AMP-CpG immunization further enabled comparable responses at 10 μg and 5 μg Spike RBD doses, and although responses at 1 μg were decreased, these still exceeded the response observed for alum (8-fold) and were similar to those generated through immunization with soluble CpG at the 10 μg dose level (Figure 8B).

**Figure 8:**
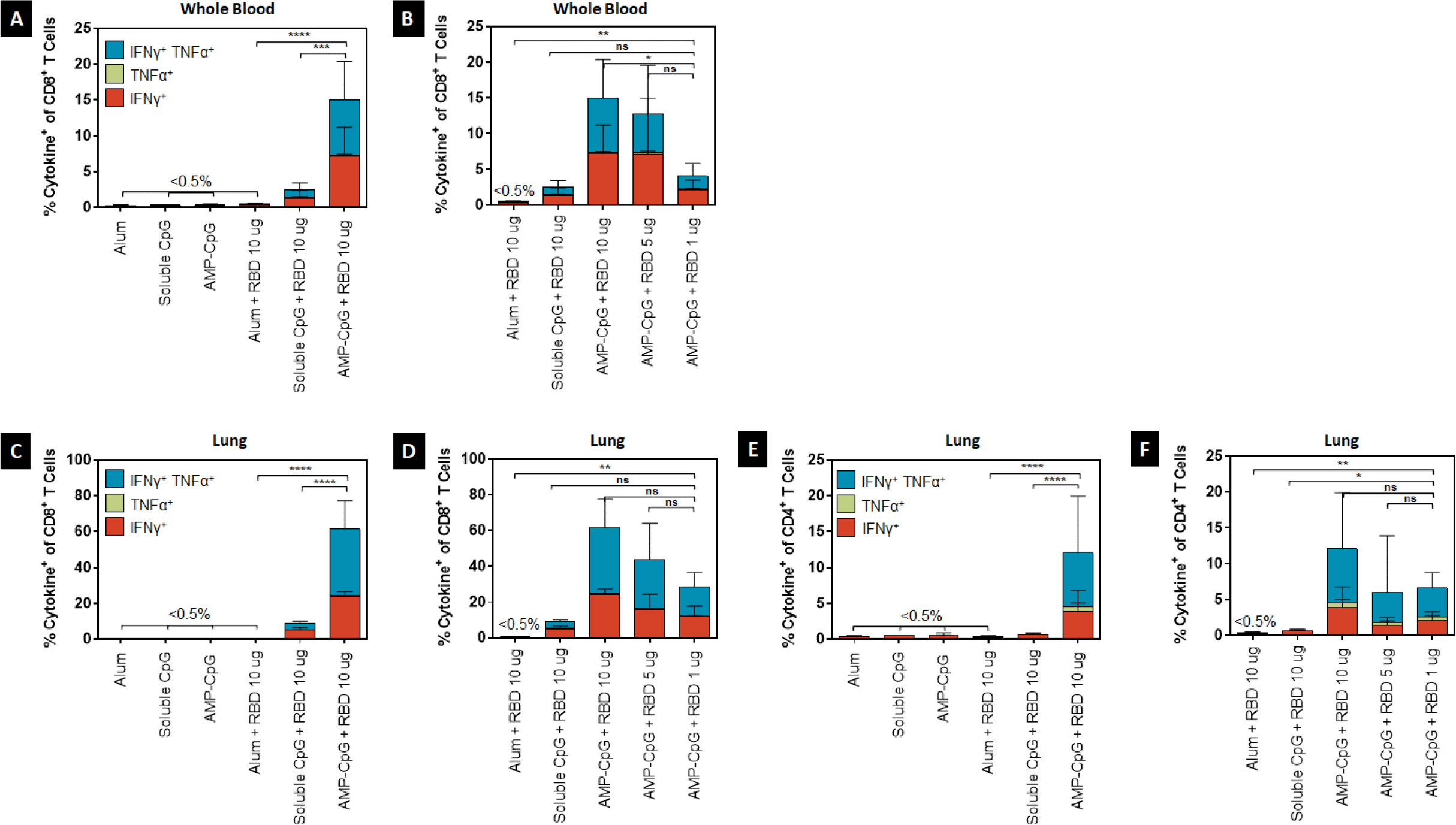
Vaccination with AMP-CpG in aged mice enables dose sparing of Spike RBD to elicit cellular immunity. 37-week-old C57BL/6J mice were immunized on day 0, 14, and 28 with 10 μg Spike RBD protein admixed with 100 μg Alum or 1 nmol soluble-CpG. Comparator animals were dosed with 1, 5, or 10 μg Spike RBD admixed with 1 nmol AMP-CpG. Control animals were dosed with adjuvant alone. T cell responses were analyzed on day 21 and 35. Cells collected from **(A-B)** peripheral blood on day 21 and **(E-H)** perfused lung tissue on day 35 were stimulated with overlapping Spike peptides and assayed for intracellular cytokine production to detect antigen-specific T cell responses. Shown are frequencies of IFNγ, TNFα, and double-positive T cells among **(A-B)** peripheral blood CD8^+^, **(C-D)** lung CD8^+^, and **(E-F)** CD4^+^ T cells. n = 10 mice per group. Values depicted are mean ± standard deviation. * P < 0.05; ** P < 0.01; *** P < 0.001; **** P < 0.0001 by two-sided Mann-Whitney test applied to cytokine^+^ T cell frequencies.

Analysis on day 35 of CD8^+^ and CD4^+^ T cells in lung tissue of aged mice showed a similar trend, with AMP-CpG immunized animals producing high frequencies of cytokine producing CD8^+^ and CD4^+^ T cells. Specifically, AMP-CpG immunization elicited Th1 cytokine production in approximately 60% of lung-resident CD8^+^ T cells, approximately 7-fold and >230-fold higher than soluble CpG and alum immunization, respectively (Figure 8C). As observed in prior studies, the elicited T cells were highly polyfunctional with more than half of the induced cells exhibiting simultaneous production of IFNγ and TNFα. Unlike the responses in peripheral blood, lung-resident cytokine producing CD8^+^ T cell frequencies did not decline in aged mice following AMP-CpG immunization relative to responses in young healthy animals and were maintained at statistically comparable levels in the 5 μg and 1 μg Spike RBD dosed groups (Figure 8D). Lung-resident CD4^+^ T cell responses exhibited a similar pattern with AMP-CpG inducing higher frequencies of Th1 cytokine producing cells (approximately 10% of CD4^+^ T cells) compared to soluble CpG (approximately 0.6% of CD4^+^ T cells) and alum (<0.5% of CD4^+^ T cells) (Figure 8E). Again, these response levels were comparable to those observed in young healthy mice showing that AMP-CpG immunization can raise comparable lung-resident T cell responses in young and aged mice. Finally, the lung-resident CD4^+^ T cell responses were maintained at comparable levels among all Spike RBD dose levels tested, and, importantly, AMP-CpG immunization at the lowest concentration of Spike RBD (1 μg) outperformed both soluble CpG and alum at a 10-fold higher antigen dose (10 μg; Figure 8F).

#### Humoral Immune Response

Spike RBD-specific antibody responses were evaluated on day 35 after repeat dose immunization with comparator vaccines in aged mice. Pseudovirus neutralization showed that AMP-CpG immunization at the 10 μg antigen dose level elicited enhanced neutralizing titers, at least 5-fold greater than those observed for soluble CpG and alum comparators, and >50-fold greater than observed in human convalescent sera/plasma (Figure 9A). Reduced doses of Spike RBD with AMP-CpG gave lower neutralizing titers which were comparable to soluble CpG and alum (Figure 9B). Of particular interest was the equivalency of titers from animals immunized with 10 μg Spike RBD with soluble CpG or alum relative to those receiving the lower 1 μg Spike RBD dose with AMP-CpG. Assessment of total IgG showed AMP-CpG and alum produced comparable Spike-RBD-specific IgG titers, both in excess of that generated in soluble CpG immunized animals (Figure 9C). Although a significant decline was observed in IgG titer with decreasing Spike RBD dose in AMP-CpG immunized animals, there was no statistical difference between AMP-CpG with 1 μg Spike RBD and alum with 10 μg Spike RBD. Isotype analysis yielded similar observations to those made in young healthy mice, with AMP-CpG driving more Th1, IgG2bc-dominant responses compared with soluble CpG or alum, which yielded more balanced or Th1, IgG1-biased profiles (Figure 9D-G). No significant difference was observed among AMP-CpG immunized animals at the varying dose levels of Spike RBD (Figure 9F), although the strength of Th1-bias observed for AMP-CpG immunized mice was reduced in aged mice relative to young healthy mice. As previously observed in young healthy mice, IgG3 titers were enhanced in AMP-CpG immunized animals compared with soluble CpG or alum (Figure 9G). Together, these results show the potential for AMP-CpG to elicit potent and functional Spike RBD-specific humoral immunity in aged mice beyond what was observed for soluble CpG or alum vaccine comparators while producing an optimal Th1-biased isotype profile and enabling at least 10-fold dose sparing of antigen.

**Figure 9:**
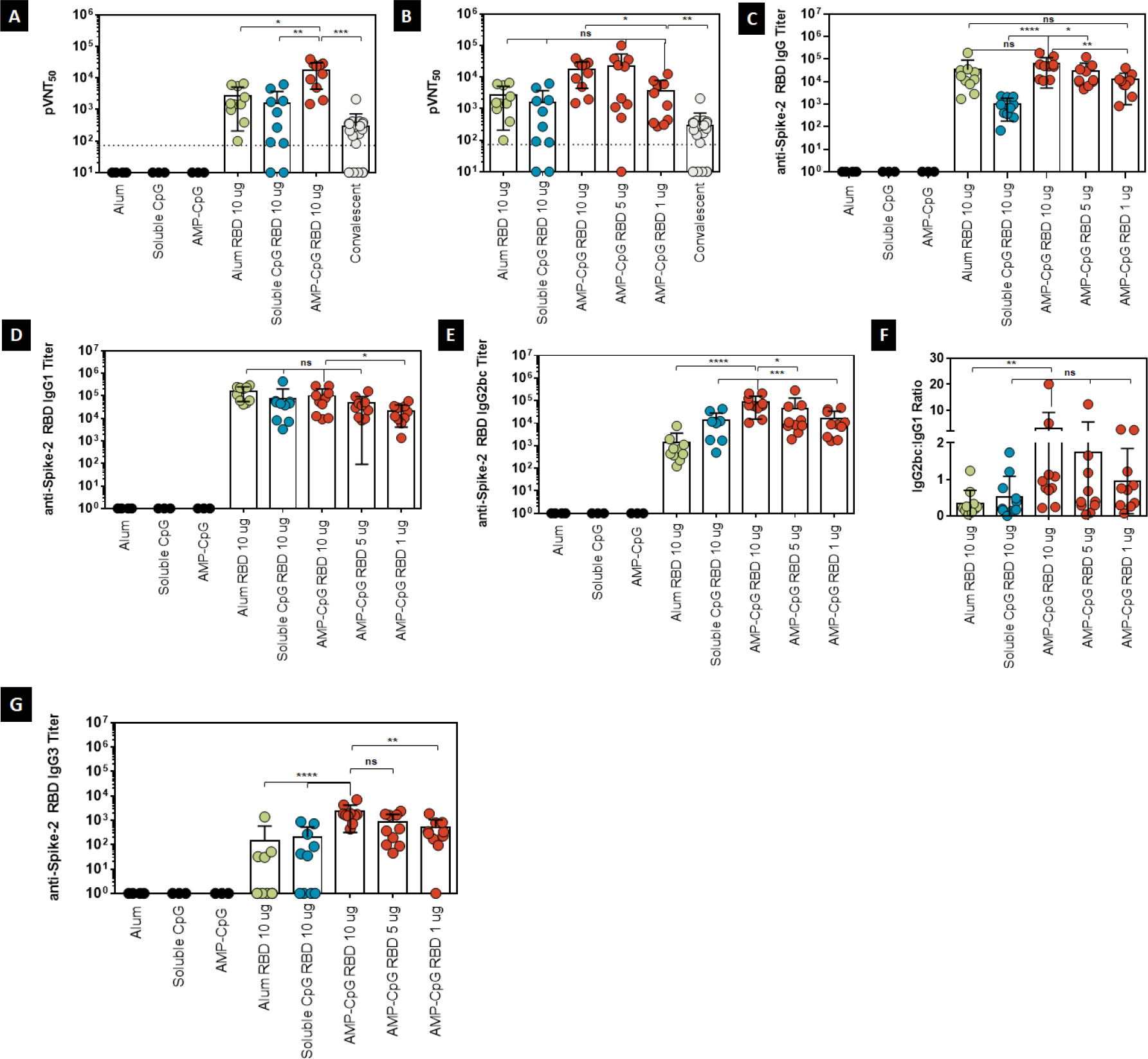
Vaccination with AMP-CpG in aged mice enables dose sparing of Spike RBD to elicit humoral immunity. 37-week-old C57BL/6J mice were immunized on days 0, 14, and 28 with 10 μg Spike RBD protein admixed with 100 μg Alum or 1 nmol soluble-CpG. Comparator animals were dosed with 1, 5, or 10 μg Spike RBD admixed with 1 nmol AMP-CpG. Control animals were dosed with adjuvant alone. Humoral responses were analyzed on day 35. Humoral responses specific to Spike RBD were assessed in serum from immunized animals by ELISA or pseudovirus neutralization assay. Shown are **(A, B)** pseudovirus neutralization ID_50_ (pVNT_50_) on day 35, **(C-G)** endpoint titers and endpoint titer ratios determined for **(C)** IgG, **(D)** IgG1, **(E)** IgG2bc, **(F)** IgG2bc:IgG1 ratio, and **(G)** IgG3 on day 35. Pseudovirus neutralization activity was compared to serum/plasma from a cohort of 22 convalescent patients who had recovered from SARS-CoV-2 infection. n = 10 mice per group. Values depicted are mean ± standard deviation. Not detected values are shown on the baseline; * P < 0.05; ** P < 0.01; *** P < 0.001; **** P < 0.0001 by two-sided Mann-Whitney test. Pseudovirus LOD (indicated by the dotted line) was determined as mean + 90% CI calculated for mock treatment.

## Discussion

Development of a safe and effective SARS-CoV-2 vaccine is urgently needed considering the profound public health, economic, and social impacts of the COVID-19 global pandemic. Multiple parallel efforts to generate a COVID-19 vaccine are ongoing to improve the chance of success^40,41^.

Emerging data from SARS-CoV-2 patients and earlier data from individuals with MERS and SARS infections, have provided information about the natural immune response to COVID-19. SARS-CoV-2 neutralizing antibody responses to infection are temporary, declining quickly after recovery^16^. In addition, recent studies have shown that a subset of patients recover from COVID-19 with SARS-CoV-2 specific T cells but not neutralizing antibodies^8,9,10^, indicating a potentially important role for T cells as a mechanism of disease prevention or mitigation. Indeed, in mouse models, CD8^+^ and CD4^+^ T cells were necessary for protective immunity against SARS and MERS infections^42,43^. Further indication of T cell mediated disease modification is evident in studies of COVID-19 patient outcomes where age-dependent reduction of T cell numbers in patients greater than 60 years of age correlated with increased COVID-19 disease severity^11^. Patients who recovered from COVID-19 without requiring intensive care had relatively high T cell levels^4,5,6,7^, versus low T cell levels observed in patients who died^11^ or who had severe disease^44^, suggesting that an increased level of T cell induction could lead to clinical benefit. Moreover, while antibody responses to SARS-CoV-2 are transient^14,15,16^, T cell responses following infection to prior betacoronaviruses, SARS and MERS, endure for decades^12^, indicating that T cells may increase the duration of protection. Several studies have now shown that the natural immune response to SARS-CoV-2 induces CD8^+^ and CD4^+^ T cells specific for epitopes within Spike RBD^34,46^.

Based on these emerging COVID-19 translational immunogenicity data, in addition to being safe, an optimal SARS-CoV-2 vaccine should **(1)** induce robust and durable CD8^+^ and CD4^+^ T cell responses, **(2)** elicit high magnitude neutralizing antibodies, **(3)** produce Th1 bias in the elicited antibody and T cell responses, **(4)** potentially expand pre-existing cross-reactive T cells, **(5)** enable dose-sparing of required immunogens to improve the speed and cost of broad vaccination campaigns, and **(6)** be efficacious in elderly populations.

Here we describe the evaluation of a novel lymph-node-targeting protein-subunit vaccine, ELI-005, comprised of AMP-CpG adjuvant paired with Spike RBD. The AMP-CpG adjuvant is designed with a key structural subunit, an albumin-binding lipid domain, that exploits the natural trafficking of albumin from subcutaneous tissue at the injection site into lymph nodes to efficiently deliver the immune activating adjuvant CpG to antigen-presenting dendritic cells^24,26^. Prior studies evaluating CpG as a vaccine adjuvant have demonstrated the importance of achieving lymphatic delivery^24,26,47,48^. While conventional unmodified CpG exhibits poor lymphatic accumulation correlated with detectable but modest induction of antigen-specific T cell responses, strategies achieving enhanced delivery of CpG directly into lymph nodes have yielded substantially stronger T cell responses 10-1000-fold greater than those observed with conventional controls^24^. Albumin-binding immunogens generated using the amphiphile strategy have proven to be among the most efficient lymph node targeting agents reported preclinically^23^, and albumin-mediated lymphatic transport has been proven effective in humans through extensive use of albumin-binding imaging agents for visualization of sentinel lymph nodes post solid tumor resection^49,50,51,52^. Thus, the application of this approach to immunization against SARS-CoV-2 is uniquely positioned to induce potent immunity through optimal lymph node targeting based on validated mechanisms suggesting effective translation to clinical use.

Generally, COVID-19 vaccine candidates have induced low T cell responses in mice, with IFNγ^+^ responses of 0-2% and 0-1% among CD8^+^ and CD4^+^ T cells, respectively^54,55,56^, or 1,000-2,000 SFC/1×10^6^ splenocytes by IFNγ ELISpot^30,31,53,55,56,57^. Consistent with these observations, a recent study evaluating immunization with soluble CpG in combination with Spike-2 protein in mice, reported no detectable cytokine production in assays of splenic T cells stimulated with Spike-2^58^. In contrast, we observed that immunization with AMP-CpG is associated with increased quantity and function in responding T cells, inducing 44% of CD8^+^ and 1.5% of CD4^+^ T cells producing IFNγ and TNFα. ELI-005-induced T cells were also highly polyfunctional, with a large fraction of antigen-specific cells producing multiple Th1 effector cytokines, consistent with the expectation that T cells receiving abundant antigen-presentation and robust costimulation from professional antigen-presenting cells in the lymph nodes acquire improved characteristics of effector function^27^. T cells generated by ELI-005 immunization trafficked in large numbers to both lung parenchyma and respiratory secretions establishing a robust T effector population ready for rapid detection and response to viral infection at a key site of first exposure. By comparison, in our studies, soluble CpG and alum immunization produced significantly weaker responses comparable to those commonly observed in other studies of preclinical vaccine candidates in mice. These results are consistent with the prior evidence demonstrating the potential for improved immunogenicity correlated with enhanced lymph node accumulation.

Antibody responses to ELI-005 showed 265-fold higher neutralization titers than sera/plasma from convalescent human COVID-19 patients. ELI-005 exceeds or matches other vaccine candidates in development of pseudovirus neutralization titers^30,31,53,54,55,56,57^. As these results compare favorably to neutralization responses (pVNT_50_ 5-40) that prevented SARS-CoV-2 pneumonia in rhesus macaques after vaccination with a chimpanzee adenoviral vector vaccine^32,56^ and to responses (pVNT_50_ 74-170) that led to rapid >3 log_10_ decreases in viral load after DNA vaccination^32^, ELI-005 holds potential to reduce disease severity as well as increase the humoral response beyond natural infection.

Previously, CD4 Th2 phenotype has been associated with vaccine associated enhanced respiratory disease in patients vaccinated against rubella^59^, respiratory syncytial virus^60^, and in preclinical studies of potential SARS and MERS vaccines^61,62^. For this reason, we characterized the spectrum of cytokines released from ELI-005 induced T cells in the lung, finding a strong Th1 skew with IFNγ and TNFα and no evidence of Th2 or Th17 cytokine secretion. As with the T cell response to ELI-005, there was a Th1 skew in the antibody response: IgG2 and IgG3 levels were increased, while IgG1 levels were generally much lower than for comparator vaccines. In mice, IgG2 isotypes (analogous to human IgG1) are the preferred IgG subclass exhibiting optimal activity against viral infections^63,64^. Specifically, murine IgG2 exhibits an enhanced ability to induce complement and antibody-dependent cell-mediated cytotoxicity via optimal interactions with an elongated and flexible Fc-region, which is consistent with a Th1 humoral response.

Through recent clinical studies, several groups^13,34,65^ have now demonstrated that patients lacking prior exposure to COVID-19 have pre-existing memory T cell responses which represent cross-reactive T cell clones cognate to common cold coronavirus epitopes. This suggests an additional potential benefit for translation of ELI-005 as immunization with AMP-CpG is expected to rapidly expand pre-existing pools of cross-reactive T cells resident in the lymph nodes, while vaccine strategies with less effective lymph node accumulation may have limited ability to access these memory cells for activation. Since the T cell immune correlates of protection in humans are not yet known for SARS-CoV-2, clinical data correlated to translational T cell immunogenicity results will be required to test this hypothesis.

As there is an urgent need to manufacture an effective vaccine for the entire global community, we examined dose sparing of the Spike RBD protein antigen and evaluated the resulting T cell and antibody responses. Even when the antigen dose was reduced from 10 μg to 1 μg, immunization with AMP-CpG maintained robust T cell responses significantly above alum and soluble CpG controls. For the humoral response, there was no significant decrease in neutralizing titer as antigen dose decreased from 10 μg to 1 μg. These results indicate that ELI-005 may have a reduced requirement for antigen dose, which could speed the delivery of protein subunit vaccines to large populations and reduce the cost and logistical burden of large-scale manufacturing efforts. Further, a protein-based vaccine with an AMP adjuvant provides the potential to develop a stable liquid formulation, simplifying distribution without frozen shipment.

Effective immunization in the elderly has proven challenging using conventional subunit vaccination due to immunological senescence. However, when soluble CpG was used as the adjuvant for the marketed hepatitis B vaccine, Heplisav-B^®^, the proportion of 60-70 year-olds who achieved seroprotection increased by 27.3% compared to alum^66^, suggesting that agonism of the toll-like receptor 9 pathway via CpG DNA adjuvants may have a unique potential for inducing improved responses in aged populations. Therefore, we investigated the potential for ELI-005 to elicit immune responses in aged mice. The results demonstrate that while neutralizing antibody titer and peripheral blood T cell responses were reduced compared to responses observed in younger mice, these responses were still significantly stronger than those induced by soluble CpG or alum comparators, and in excess of levels observed in convalescent patients. Combined with the evidence that CpG is an effective adjuvant for immunization in the elderly, these results suggest that ELI-005 should be evaluated in nursing home patients who have had the highest COVID-19 mortality rates during the current pandemic.

Further research will be needed to assess the potential for AMP-CpG to increase the duration of protective neutralizing antibody and functional T cell responses. Though our data assessed responses to SARS-CoV-2, the general features of the T cell response recapitulated the prior observations of AMP vaccination^25,26^ in oncology, while demonstrating parallel potential to induce neutralizing antibody responses. There is potential for AMP-CpG to enhance a broad spectrum of novel vaccines. The promising immunogenicity assessments reported here for ELI-005 support its clinical evaluation as a potential vaccine for SARS-CoV-2.

## Methods

### Animals

Female, 6 to 8-week-old C57BL/6J and BALB/c mice and 37-week-old C57BL/6J mice were purchased from Jackson Laboratory (Bar Harbor, ME). All animal studies were carried out under an institute-approved IACUC protocol following federal, state, and local guidelines for the care and use of animals. Mice were injected with 1 nmol CpG (soluble CpG), 1 nmol lipid-conjugated CpG (AMP-CpG), or 100 μg Alum admixed with phosphate-buffered saline (PBS) only (adjuvant controls), or 1-10 μg of Spike RBD protein (Sino Biological, Cat: 40592-V08H or GenScript, Cat: Z03483). “Mock” groups received PBS alone. Injections (100 μL) were administered subcutaneously at the base of the tail (50 μL bilaterally) on days 0, 14, and 28. Blood samples were collected on days 7, 21, and 35. Mice were sacrificed on day 35 for lung harvest and collection of bronchoalveolar lavage (BAL) fluid. Only the pilot experiment (Figure 1) received a fourth dose on day 42 and samples were collected on day 49. Lungs were harvested following perfusion with 10 mL of PBS into the right ventricle of the heart. Lung tissue was physically dissociated and digested with RMPI1640 media containing 1 mg/mL collagenase D and 25 units/mL DNaseI. BAL fluid was obtained by washing the lungs three times each with 1 mL PBS.

### Human Convalescent Serum/Plasma

Convalescent serum samples (n=7) and plasma samples (n=15) from patients who had recovered from SARS-CoV-2 infection (COVID-19) were obtained from US Biolab (Rockville, MD) and ALLCELLS (Alameda, CA), respectively. All samples were received and stored frozen at -80°C until analysis.

### Antigen-Binding ELISA

ELISAs were performed to determine sera antibody binding titers. ELISA plates were coated with 200 ng/well CoV2 RBD protein (GenScript; Cat: Z03483) overnight at 4°C. Plates were pre-blocked with 2% bovine serum albumin (BSA) for 2 h at room temperature (RT). Serially diluted mouse sera were transferred to the ELISA plates and incubated for 2 h at RT. Plates were washed 4 times with washing buffer (BioLegend; Cat: 4211601) and then incubated for 1 h at RT with a 1:2000 dilution of secondary antibody of horse radish peroxidase (HRP)-conjugated rabbit anti-mouse IgM (μ chain), HRP-rabbit anti-mouse IgG (Fcγ), HRP-goat anti-mouse IgG1, HRP- goat anti-mouse IgG2b, HRP- goat anti-mouse IgG2c, or HRP-goat anti-mouse IgG3 (Jackson ImmunoResearch, Cat: 315-035-048, 315-035-049, 315-035-046, 115-035-205, 115-035-207, 115-035-208, and 115-035-209, respectively). Plates were again washed 4 times with washing buffer; after which the plates were developed with 3,3’,5,5’-tetramethytlbenzidine for 10 min at RT, and the reaction was stopped with 1N sulfuric acid. The absorbance at 450 nm was measured by an ELISA plate reader. Titers were determine at an absorbance cutoff of 0.5 OD.

### SARS-CoV-2 Pseudovirus Neutralization Assay

For the neutralization assay we used the ACE2-HEK293 recombinant cell line (BPS Bioscience, Cat: 79951) or the control HEK293 cell line (ATCC) and the SARS-CoV2 Spike Pseudotyped Lentivirus (BPS Bioscience, Cat: 79942). The SARS-CoV2 Spike Pseudotyped Lentivirus contains the luciferase reporter gene and the SARS-CoV2 Spike envelope glycoproteins, thus specifically transducing ACE2-expressing cells. Mouse or human sera dilutions were performed in the Thaw Medium 1 (BPS Bioscience, Cat: 60187) in 96-well white clear-bottom luminescence plates (Corning, Cat: 3610) and then pre-incubated with 10 µL of virus for 30 minutes at RT. ACE2-HEK293 or control HEK293 cells (40 µL), containing 10,000 cells, were then added to the wells and incubated at 37°C for 48 h. Control wells included ACE2-HEK293 cells or control HEK293 cells with the virus, but no sera, and provided the maximum transduction level and the background, respectively. Luciferase activity was detected by adding 70 µL of freshly prepared ONE-Step Luciferase reagent (BPS Bioscience, Cat: 60690) for 15 minutes at RT and luminescence was measured with a Synergy H1 Hybrid reader (BioTek). Pseudovirus neutralization data for the pilot experiment was performed by GenScript (Nanjing, China) following the same protocol, but using in-house ACE2-HEK293 cells and Spike RBD-HRP recombinant protein. Pseudovirus neutralization titers at the half-maximal inhibitory dilution (pVNT_50_) were calculated as the serum dilution at which RLU were reduced by 50% compared to RLU in virus control wells.

### SARS-CoV-2 Surrogate Neutralization Assay

For the surrogate neutralization assay the cPass kit from GenScript was used (Cat: L00847). Manufacturer’s instructions were followed. In short, serially diluted sera were incubated with SARS-CoV-2 Spike RBD-HRP and added to ACE2 precoated plates. Plates were developed with 3,3’,5,5’-tetramethytlbenzidine for 10 min at RT, and the reaction was stopped with 1N sulfuric acid. The absorbance at 450 nm was measured by an ELISA plate reader. Titers were determined at an absorbance cutoff of 0.5 OD.

### Lung-Resident T Cell Cytokine Determination by Cytometric Bead Array (CBA)

CBA flow cytometry was performed to determine cytokine production. Lung-resident leukocytes (collected after the final booster dose) were activated overnight with SARS-CoV-2 Spike glycoprotein overlapping peptides at 1 μg/peptide per well (consisting of 315 peptides, derived from a peptide scan [15-mers with 11 amino acid overlap] through Spike glycoprotein of SARS-CoV-2) (JPT, Cat: PM-WCPV-5 or GenScript, Cat: RP30020). Phorbol Myristate Acetate (PMA, 50 ng/mL) and ionomycin (1 μM) were used as positive controls, and complete medium only as the negative control. Culture supernatants were harvested, and Th1/Th2 cytokine production was measured (CBA Mouse Th1/Th2/Th17 Cytokine Kit: BD, Cat: BDB560485). Briefly, bead populations with distinct fluorescence intensities that are coated with capture antibodies specific for various cytokines including IFNγ, TNFα, IL-4, IL-6, IL-10, and IL-17 were incubated with culture supernatants. The different cytokines in the sample were captured by their corresponding beads. The cytokine-captured beads were then mixed with phycoerythrin (PE)-conjugated detection antibodies. Following incubation, samples were washed, and fluorescent intensity of PE on the beads were measured and analyzed by flow cytometry (BD FACSCanto II). Mean fluorescent intensities (MFI) were calculated using FACSDiva software (BD) and protein concentrations were extrapolated using Microsoft Excel.

### Peripheral Blood and Lung-Resident T Cell Cytokine Determination by Intracellular Cytokine Staining (ICS)

ICS was performed for TNFα and IFNγ. Peripheral blood cells (collected 7 days after each booster dose) and lung-resident leukocytes (collected after the final booster dose) were stimulated overnight with 1 μg/peptide per well of Spike-derived overlapping peptides at 37°C, 5% CO_2_ in the presence of brefeldin A (Invitrogen, Cat: 00-4506-15) and monensin (BioLegend, Cat: 420701). Cells were stained with the following antibodies: PE anti-mouse IFNγ (BD, Cat: 554412), FITC anti-mouse TNFα (BD, Cat: 554418), APC-Cy^™^7 anti-mouse CD3 (BD, Cat: 560590), PE-Cy7 anti-mouse CD4^+^ (Invitrogen, Cat: 25-0041-82), and APC anti-mouse CD8a (eBioscience, Cat: 17-0081-83). PMA (50 ng/mL) and ionomycin (1 μM) were used as positive controls, and complete medium only as the negative control. Cells were permeabilized and fixed (Invitrogen, Cat: 00-5523-00). A LIVE/DEAD fixable (aqua) dead cell stain kit (Invitrogen, Cat: L34966) was used to evaluate viability of the cells during flow cytometry. Sample acquisition was performed on FACSCanto II (BD) and data analyzed with FlowJo V10 software (TreeStar).

### IFNγ ELISpot

Spleens from mice were collected individually in RPMI 1640 media supplemented with 10% FBS and penicillin, streptomycin, nonessential amino acids, sodium pyruvate, and beta-mercaptoethanol (complete media) then processed into single cell suspensions and passed through a 70 μm nylon filter. Cell pellets were re-suspended in 3 mL of ACK lysis buffer (Quality Biological, Inc., Cat: 118156101) for 5 min on ice; then PBS was added to stop the reaction. The samples were centrifuged at 400×g for 5 min at 4°C and cell pellets were re-suspended in complete media. ELISpot assays were performed using the Mouse IFN-γ ELISpot Set (BD, Cat: BD551083). 96-well ELISpot plates precoated with capture antibody overnight at 4°C were blocked with complete media for 2 h at RT. 500,000 mouse splenocytes were plated into each well and stimulated overnight with 1 μg/peptide per well of Spike-derived overlapping peptides. The spots were developed based on manufacturer’s instructions. PMA (50 ng/mL) and ionomycin (1 μM) were used as positive controls, and complete medium only as the negative control. Spots were scanned and quantified by an ImmunoSpot CTL reader.

### Statistics

All data were plotted and all statistical analyses were performed using GraphPad Prism 8 software (La Jolla, CA). All graphs display mean values and the error bars represent the standard deviation. No samples or animals were excluded from the analyses. Animals were not randomized for any of the studies, and dosing was not blinded. Statistical comparisons between groups were conducted using two-sided Mann-Whitney tests. Data were considered statistically significant if the p-value was less than 0.05.

## Acknowledgements

The authors would like to thank Deborah M. Lidgate for expert medical writing assistance, Charles S. Davis at CSD Biostatistics, Inc., Julian Adams at Gamida Cell, Inc., and Darrell Irvine at M.I.T. for helpful advice and discussion.

## Author Contributions

P.C.D., M.P.S., L.K.M, and C.M.H. designed experiments, analyzed the data, and prepared the manuscript. M.P.S., L.M.S., A.J., and L.K.M. designed and performed experiments. C.M.H. and P.C.D. provided supervision and oversaw final manuscript preparation. All authors reviewed and approved the version for publication.

## Competing Interests

All authors are employees of Elicio Therapeutics, and as such receive salary and benefits, including ownership of stock and stock options from the company. P.C.D., M.P.S., L.M.S., and C.M.H have an amphiphile SARS-CoV-2 vaccine patent pending to Elicio.

## Additional Information

Correspondence and requests for materials should be made to C.M.H.

